# *Azadirachta indica:* derived, red-emitting, carbon nanoparticles for cellular bioimaging and potential therapeutics

**DOI:** 10.1101/2024.02.26.582094

**Authors:** Stuti Gandhi, Sweny Jain, Dhiraj Bhatia, Pankaj Yadav

## Abstract

Red-emitting carbon nanoparticles (CNPs) were synthesized by the refluxed green synthesis method using ethanolic extract of neem leaves (*Azadirachta indica*). These nanoparticles were called as nQDs (neem quantum dots). The nQDs exhibited excellent photoluminescence properties with a maximum emission at 672nm, and the average size of nQDs was around 47nm. In the *in-vitro* study, Retinal Pigment epithelial (RPE1) cells and SUM159A cells showed enhanced cellular uptake. In RPE1 cells, the cellular uptake was higher than in SUM159A cells. In the biocompatibility assay, SUM159A cell viability declined with the increasing nQDs concentration. The results show that red-emissive CNPs can be synthesized from *Azadirachta indica* (neem) leaves using a simple method with a possible application in bioimaging and therapeutics.

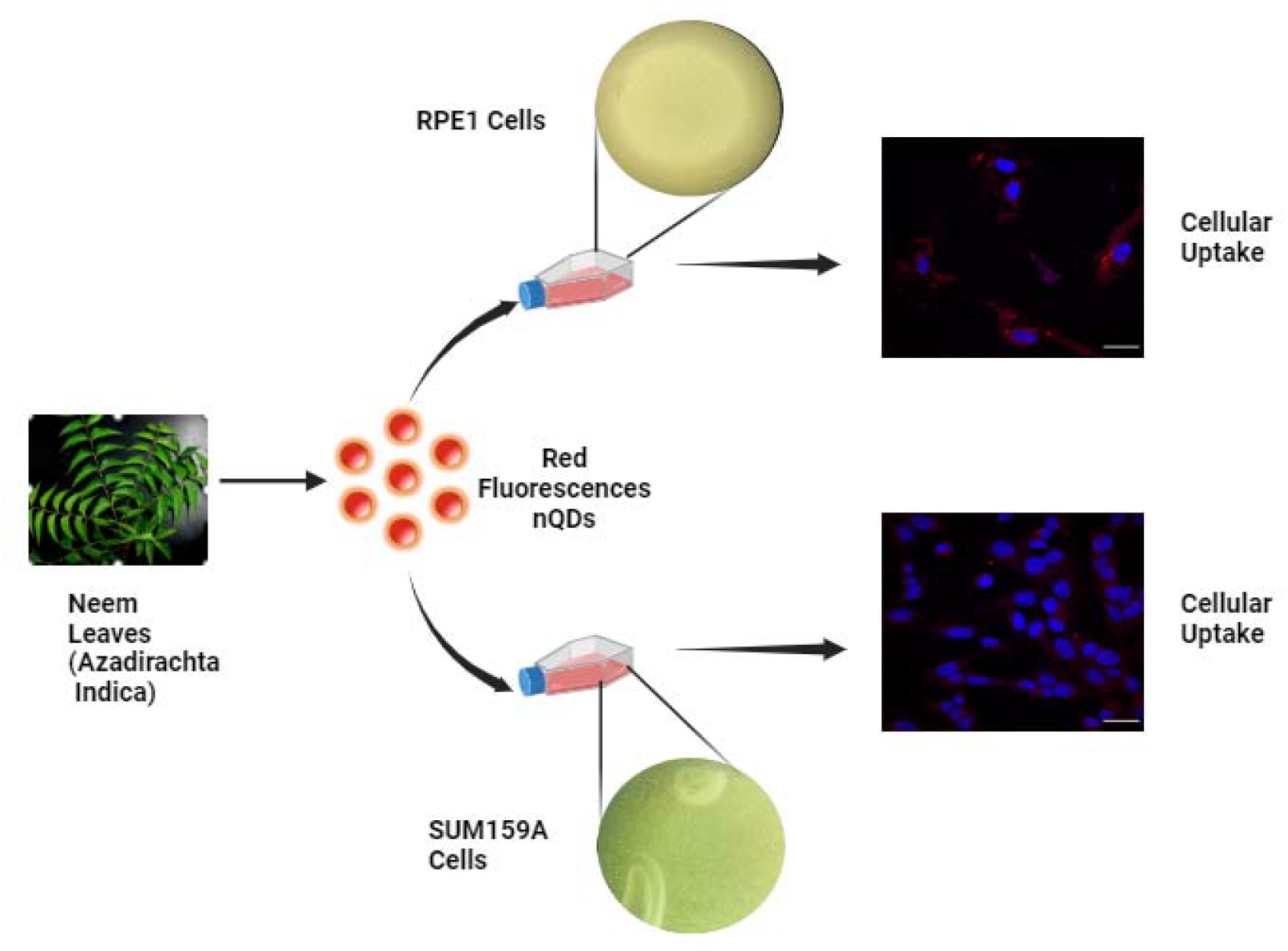

## 1. Introduction

Nanomaterials (NMs) in the 1-100 nm range have attracted significant scientific interest in the last decade and these particles are known as nanoparticles (NPs).^1^ Due to their magnetic, catalytic, mechanical, optical, and electronic properties^2^, NMs have drawn much attention in the current field of study and have numerous applications in medicine^3^, cosmetics, packaging, nanofibers, biosensors, and electronics.^4^ NMs are categorized according to their size, shape, and morphology.^1^ Quantum dots, also known as semiconductor nanoparticles, are smaller than the exciton Bohar radius^5^ and have a 1 to 10 nm diameter.^6^ They are known for their unique optoelectronic and physiochemical properties and can be utilized to research a variety of biological processes.^7^ Louis Brus discovered quantum dots (cadmium sulfide) in colloidal suspension.^8^ At the same time, Alexey Ekimov found quantum dots in a glass matrix.^9^ In 2004, Xu et al. produced carbon nanoparticles (CNPs) for the first time in history by purifying single-walled carbon nanotubes.^10^ Sun et al. dubbed these CNPs “Carbon Quantum Dots” in 2006.^11^

Carbon quantum dots (CQDs), or carbon dots (CDs), are a kind of nanomaterial with zero dimensions.^12^ CQDs are a type of carbon nanomaterial that is environmentally beneficial; they have stable physiochemical characteristics and strong biocompatibility.^13–15^ These carbon-based nanomaterials, which include graphene oxide, fullerenes, carbon nanotubes, and carbon nanodots, are extensively employed in biomedical devices, including optical sensors, diagnostic tools, and bioimaging probes that provide chemical stability and size-dependent photoluminescence.^16^ For the synthesis of fluorescent carbon nanoparticles (CNPs), a variety of precursor materials are employed; however, due to their affordability, environmental friendliness, and tuneable fluorescent CNPs, the uses of CNPs in bioimaging, sensing drug delivery, catalysis, photovoltaics, and other fields are expanding.^17^ The fluorescence characteristics of CNPs under ultraviolet (UV) irradiation are their most distinctive features.

Two methods are used for synthesizing CNPs: the Top-down and Bottom-up methods. The top-down process uses arc discharge, electrochemical, oxidation, and laser ablation to create CNPs from massive graphitic materials. However, they need costly materials, time-consuming procedures, and harsh reaction conditions.^18,19^ The bottom-up strategy uses pyrolysis, microwave irradiation, reflux method, and hydrothermal/solvothermal synthesis to create CNPs from small molecules. The bottom-up method is more economical and environmentally friendly, making it more suitable for CNP production.^20^ The green synthesis method is part of the bottom-up method. Natural resources and biomass, such as vegetables, plant leaves, fruits, and human derivatives, are used to synthesize green carbon nanoparticles or CNPs. The exclusive benefits are chemical-free synthesis, easy conjugation with biomolecules with small particle sizes, tuneable photoluminescence (PL), and suitable operating conditions.^21^ In 2012, the hydrothermal treatment of orange juice was used to synthesize the carbon dots.^22^ Carbon quantum dots are synthesized by strawberries and apples, which show blue fluorescence under 365nm in UV light in 2013 and 2014, respectively.^23,24^ Red emissive CNPs derived from spinach leaf powder at an excitation wavelength of 672nm.^7^ Psidium guajava (Guava) was used to derive red fluorescence carbon nanoparticles (CNPs) by microwave-assisted method.^17^ The rose-heart radish showed blue luminescence under UV light at 365nm.^25^ The three fluorescent carbon nanoparticles (FCP) were synthesized by mango fruit.^26^

We used the neem plant to synthesize the carbon nanoparticles (CNPs). The *Azadirachta indica*(Neem) is a multipurpose medicinal shrub. Neem has various applications, including anti-inflammatory, anti-cancer, anti-fungal, and anti-viral.^27^ Since 2014, hydrothermal treatment has been used to synthesize graphene quantum dots using Neem leaves; they were red at 600 nm. The average size and quantum yield (QY) of graphene quantum dots were 5 nm and 41.2%, respectively. Using the red-green-blue (RGB) color mixing technique, the N-GQDs (GQDs from neem) were used to create a white light conversion cap.^28^ Green emissive carbon nanoparticles (CNPs) were synthesized by *Azadirachta indica*(Neem) in 2019. This method was time-consuming. In this paper, the cell imaging and phototherapy of HeLa cancer cells and normal NIH-3T3 cells were performed.^29^ The fluorescent N-CQDs were synthesized from neem leaves, which showed excitation-dependent fluorescence emissions in a range of 290-370nm in 2018, dubbed by Pradeep Kumar Yadav. The one-pot hydrothermal method was used to synthesize the N-CQDs. The Quantum Yield (QY) of N-CQDs was 27.2%. In this paper, the N-CQDs were observed in Michaelis-Menten Kinetic activity and steady-state kinetics. ^30^ In another study, the microwave method was used for synthesizing red-emissive CNPs named NNPs, which were derived from neem leaves. These NNPs showed antioxidant properties at two different concentrations.^21^ In all the above studies reported, the synthesis methods were not time-effective, eco-friendly, and cost-effective. Moreover, in all the reported studies the synthesized CNPs were blue and green fluorescent.

We present a time-effective, eco-friendly, and cost-effective bottom-up method to synthesize the red-emissive carbon nanoparticles (CNPs) using neem leaves. The reflux method was used to produce nQDs that emit bright red fluorescence. This is the first time using the reflux method to synthesize carbon nanoparticles (CNPs) using neem leaves (*Azadirachta indica*). The nQDs are characterized by DLS (dynamic light scattering), FTIR (Fourier transform infrared spectroscopy), AFM microscopy for topology and morphology, UV-vis absorbance, and UV fluorescence spectroscopy for measuring fluorescence intensity. Mammalian cell lines were used to study nQDs. We used two cell lines: 1. RPE1 cells (non-transformed alternative to cancer cell line) and 2. SUM159A cells (triple-negative breast cancer cell line). These cell lines were used to study biocompatibility and cellular uptake of nQDs.

## 2. Materials and Methods

The fresh Neem leaves (*Azadirachta indica*) were collected from the IIT Gandhinagar campus. Ethanol (>99.9%) was purchased from Changshu Hongsheng Fine Chemicals Co. Ltd. The deionized water used in the experiment was taken from Merck Millipore, and a filter of 0.22 μm was purchased from Merck. The cell culture dishes, Dimethyl Sulfoxide (DMSO), and Paraformaldehyde were purchased from Himedia. Phalloidin and MTT [3-(4,5-Dimethylthiazol-2-yl)-2,5-diphenyltetrazolium bromide) were purchased from Sigma Aldrich. DMEM (Dulbecco’s Modified Eagle’s Medium), Ham’s F12 media, FBS (fetal bovine serum), and trypsin-EDTA (0.25%) from Gibco. No additional sterilization or treatment is required because all the substances were of good scientific quality.

### 2.1. Synthesis of nQDs

The leaves of *Azadirachta indica*(Neem leaves) were collected/obtained from the IIT Gandhinagar campus. The leaves were washed with Milli-Q Water to remove the dust and then allowed to air dry in the shade. The leaves were then ground into a powder using a household grinder. After that, the powder was dissolved in ethanol in a 1:10 (W/V) ratio and constantly stirred (at 300 rpm) for four hours with a magnetic stirrer. The dissolved powder is centrifuged at 10,000 rpm for 10 minutes at room temperature to extract the neem leaf extract. To produce nQDs, the extract is refluxed for two hours at 16°C and then cooled naturally. Following formation, the nQDs were filtered via a 0.22 μm syringe filter. Lastly, rotavapor was used to evaporate the solvent and produce powdered nQDs. The nQDs powder is then characterized and further used to study their cellular uptake and tissue bioimaging.

### 2.2. Characterization of nQDs

Spectrocord-210 Plus Analytokjena (Germany) and FP-8300 Jasco spectrophotometer (Japan) were used to measure the optical properties of nQDs, including both UV-Vis absorbance and fluorescence emission, respectively. A freshly peeled mica sheet was used for the AFM sample preparation. A 5μL drop of powdered nQDs (100mg/mL in water) was placed on a mica sheet. After that, the mica sheet was placed in a desiccator to dry. Lastly, the Bruker AFM equipment performed AFM imaging in tapping mode. Using Spectrum Two PerkinElmer in ATR mode, the FTIR spectra of powdered nQDs and neem leaves are recorded between 400 and 4000 cm^_1^. The Malvern analytical Zetasizer Nano ZS was used for the solution-based size characterization of nQDs by Dynamic Light Scattering (DLS). Gaussian Fit was then used to plot the data in OriginPro Software.

### 2.3. *In vitro* study

#### 2.3.1. Biocompatibility Assay

The cytotoxicity of nQDs was assessed by seeding 96-well plates with approximately 1×10^5^/100μl of SUM159A and RPE1 (Retinal Pigment Epithelium Cell Line). The plates were prepared in the following ways: Ham’s F12 complete media for SUM159A and DMEM containing 10% FBS and 1x antibiotic (pen strep) for REP1, respectively. Before the experiment, the cells were given a 24-hour acclimatization period in 5% CO_2_ at 37°C. The following day, cells were washed with PBS and treated with nQDs at increasing concentrations (50, 100, 200, 300, 400, 500μg/ml) for 24 hours in serum-free media. The next day, culture media was discarded, 100μl of media (Ham’s F12 Serum-free Media for SUM159A and DMEM Serum-free Media for RPE1) containing MTT (5mg/ml) was added, and the cells were incubated for 3-4 hours to measure the viability of cells. Later media was aspirated and 100μl of DMSO was added to each well to dissolve the purple formazan crystals. A Multiskan microplate reader was then used to measure the absorbance spectra at 570nm. The experiment was conducted in triplicate, normalized to the corresponding DMSO-containing wells, and the percentage of cell survival in each well was calculated using the non-treated nQDs well as the control.

Cells Viability (%) = OD (Sample) – OD (Blank) / OD (Control) – OD (Blank) × 100

#### 2.3.2. Cellular Uptake

The cellular uptake of nQDs was studied in the mammalian cell line; here, we used RPE1 and SUM159A cells. In a T25 flask, the SUM159A cells were cultured and maintained using Ham’s F 12 complete media. RPE1 cells were similarly cultured in a T25 flask using a DMEM complete media. Twenty-four hours before the experiment, 1 × 10^5^ cells were seeded per well on a 10mm glass coverslip on a 4-well plate. Before the experiment, cells were examined under a microscope to observe how they adhered to one another and spread. The next day, the media was removed and 450μl of serum-free media (Ham’s serum-free media for SUM159A and DMEM serum-free media for RPE1) was added, then incubated for 15 minutes at 37°C. After incubation, discard the serum-free media, and increase the concentration of nQDs (0 μg/mL, 50 μg/mL, 100 μg/mL, 200 μg/mL) were treated in cells. The cells were incubated at 37°C for 15 minutes following the treatment. The extra surface-bound nQDs were removed by washing them three times with 1X PBS For 15 min at 37°C; the cells were fixed with 4% PFA (paraformaldehyde). One more time, the fixed cells were washed three times using 1X PBS. Lastly, the coverslips were mounted on a glass slide with a Hoechst-containing mowiol to stain the nucleus. A confocal laser scanning microscope was used to take the images of the fixed cells at a resolution of 63x. Fiji ImageJ software was used to quantify the acquired pictures.

## 3. Result and Discussion

### 3.1. Synthesis and characterization of nQDs

Red fluorescent carbon nanoparticles were synthesized via the reflux method using an ethanolic extract of neem powder. Two grams of neem powder was mixed with 20 ml ethanol solution and stirred for 4 hours on a magnetic stirrer. After stirring, the mixture was centrifuged at 25°C for 10 minutes at 10,000 rpm to obtain the neem leaf extract. The supernatant was collected and refluxed at 160°C for 2 hours and cooled at room temperature. After cooling, the nQDs solution was filtered using a 0.22 μm syringe filter. The nQDs appeared green under visible light and red under UV light. Next, rotavapor was performed to evaporate the ethanol and obtain a powder of nQDs. The obtained nQDs powder was then used for further characterization and experiments.

To study the characteristics of nQDs, we used Ultraviolet-Visible (UV-Vis) absorbance spectroscopy to measure their absorbance. We used ethanol and water solvents to measure nQDs absorbance and fluorescence. Firstly, nQDs were dissolved in ethanol, for which the absorbance spectra show peaks at 267nm, 365nm, 410nm, 442nm,468nm, and 667nm, as shown in **Figure 2(a)**. The peak at 267nm represents the n-*π*\* electron transition in the carbon structure, although the peak at 354nm represents the surface or molecule center of the nQDs.^31^ Due to the energy level transition from the conjugated *π* structure of angstrom size found in nQDs, the absorbance peak was attained at a wavelength of around 442nm. The peak at 672nm is assumed to result from chlorophyll found in green plants. In water, nQDs showed distinct peaks at 267nm, 365nm, 420nm, and 667nm, as shown in **Figure 2(b)**. The peaks at 267nm, 365nm, and 667nm peaks correspond to those in water. The hydrodynamic size and zeta potential were determined for nQDs which were 295nm and -4.65mV respectively, as shown in **Figure 2(c). Figure 2(d)** displays the Fourier transforms infrared (FTIR) spectra of nQDs used to identify the functional groups and their chemical composition. N, H, C, and O atoms usually form chemical bonds on carbon-based nanoparticles. The FTIR spectra showed the presence of the following surface groups on nQDs: 3367 cm^-1^ for N-H, NH_2_, and alcohol O-H stretching, 2927 cm^-1^ for sp^3^ C-H and carboxylic acid O-H stretching, 2857 cm^-1^ for sp^3^ C-H, aldehyde C-H, carboxylic acid C-O stretching, 1734 cm^-1^ for C=O stretch, 1627 cm^-1^ for C=O, C=N, C=C alkene, and aromatic, N-H bend, 1377 cm^-1^ for sp^3^ C-H bend, 1247cm^-1^ for acyl C-O, 1161 cm^-1^ for acyl C-O stretching, 1027 cm^-1^ for alkoxy C-O stretching and 873 cm-1 for alkene sp^2^ C-H bend. The comparison of neem leaves peaks with nQDs showed peaks at 3297 cm^-1^ for sp C-H, alcohol O-H, carboxylic acid C-O, N-H_2_, and N-H stretching, 1733 cm^-1^ for C=O stretching, 1607 cm^-1^ for C=O, C=N, C=C alkene and aromatic N-H bend, 1165 cm^-1^ for acyl C-O stretching, 1107 cm^-1^ and 1037 cm^-1^ for alkoxy C-O stretching, with nQDs shows the shift of peaks in nQDs at 3367 cm^-1^, 1734 cm^-1^, 1627 cm^-1^, 1247 cm^-1^, 1161 cm^-1^ and 1027 cm^-1^. This comparison also reveals the formation of new bonds in nQDs at 1377 cm^-1^ and 873 cm^-1^.

**Figure 1.**
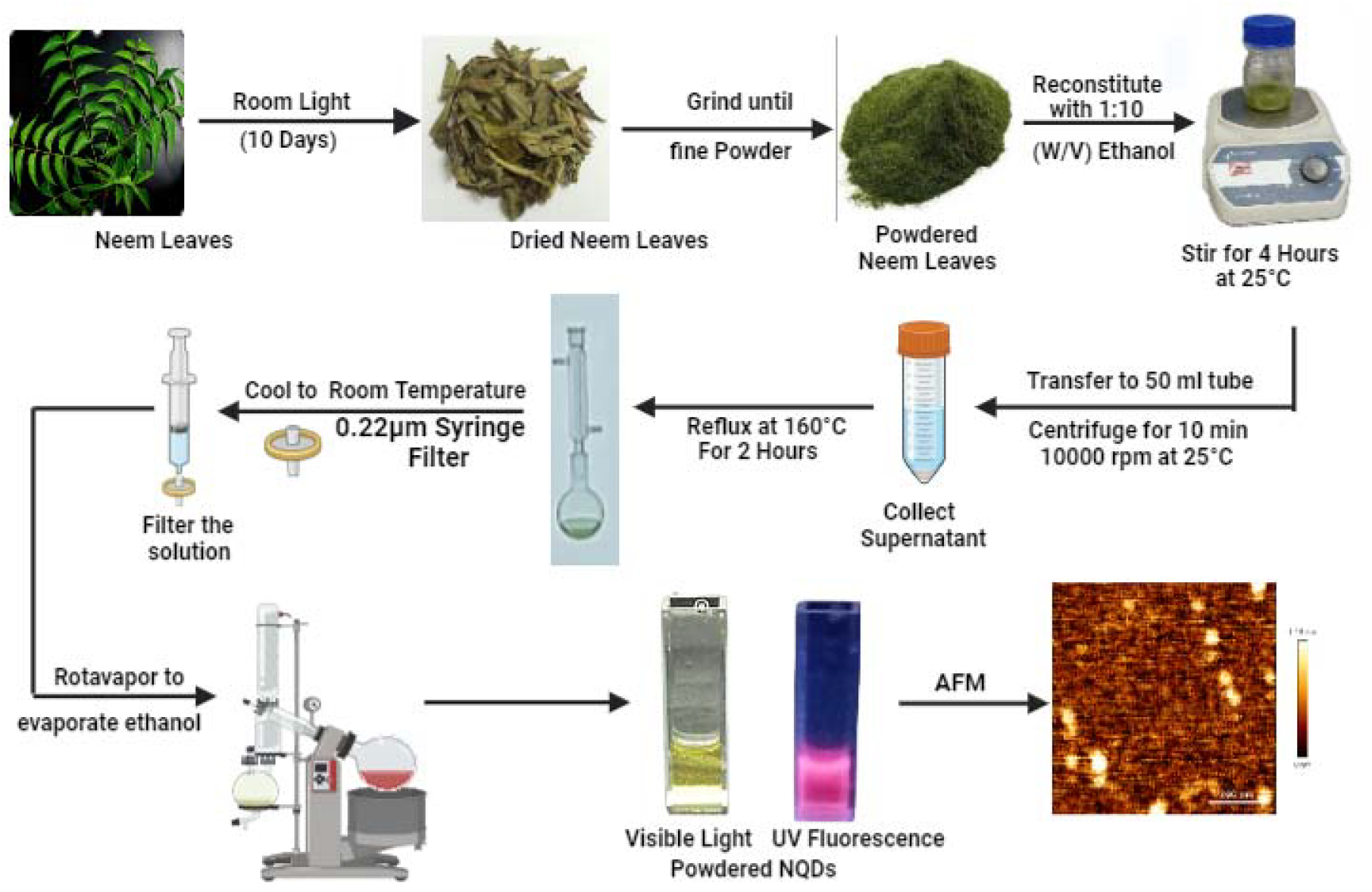
Neem leaves were used in the reflux method and synthesis of red-emitting carbon nanoparticles. The neem leaves were dried under the shade and ground to obtain neem powder. The neem powder was reconstituted with 1:10 (W/V) ethanol and stirred. The supernatant neem extract was refluxed and cooled at room temperature. Filter the neem extract, and lastly, rotavapor was used to evaporate the ethanol. After that, the nQDs were observed in visible light and UV light.

**Figure 2.**
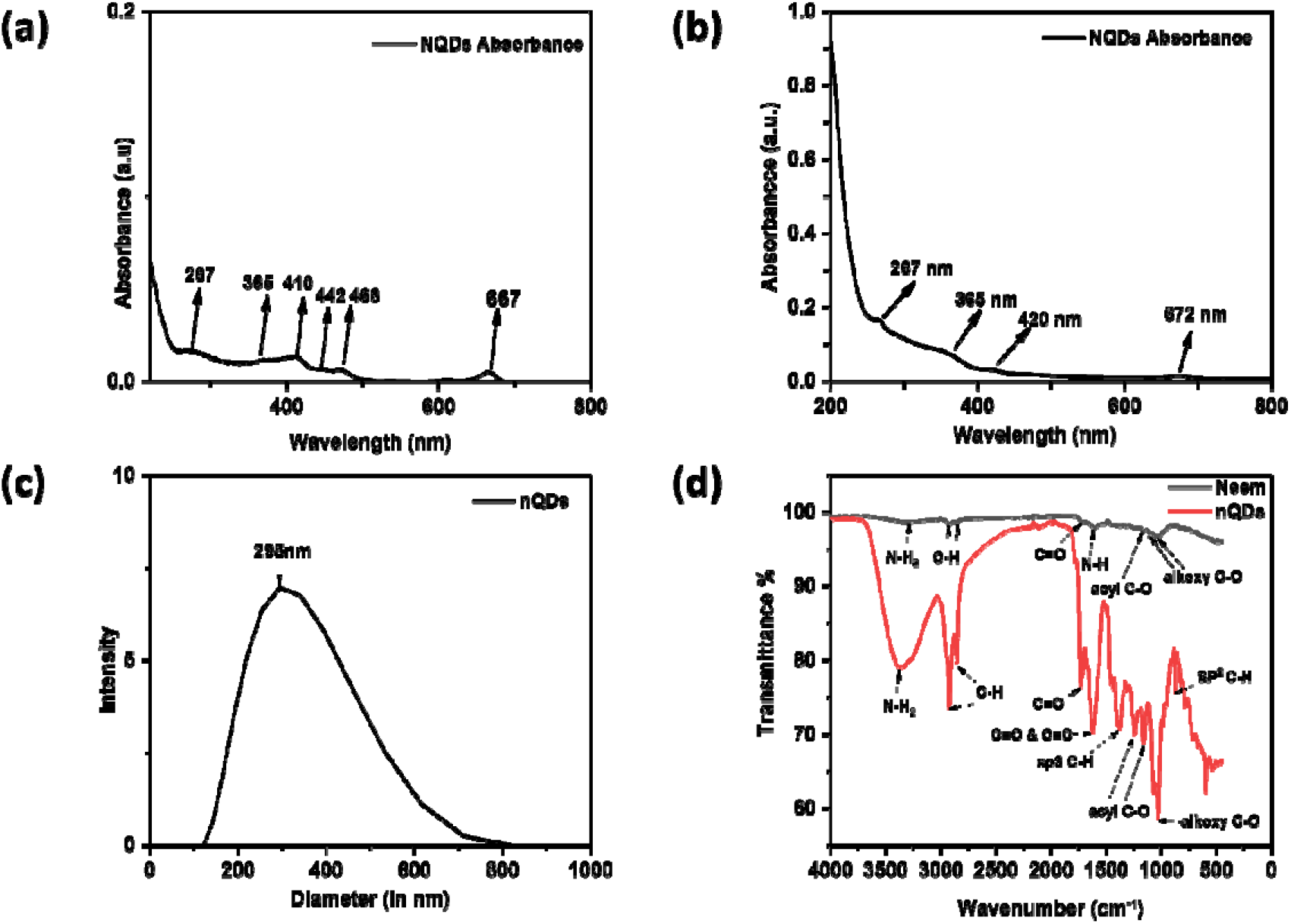
Characterization of nQDs. **(a)** The graph illustrates nQDs’ ultraviolet-visible (UV) spectra with absorbance at distinct peaks. The peaks were 267nm, 365nm, 410nm, 442nm, 468nm, and 667nm at three μg/mL in ethanol. **(b)** Ultraviolet-visible (UV) spectra of nQDs showing absorbance peaks. The peaks were 265nm, 365nm, 420nm, and 667nm at six μg/mL in water. **(c)** The hydrodynamic size of nQDs in water as a solvent was measured to be 295nm. **(d)** The chemical composition and functional groups of nQDs can be identified by their FTIR spectra. FTIR shows the surface groups on nQDs: 3367 cm^-1^ for N-H_2_, 2927 cm^-1^ and 2857 cm^-1^ for C-H, 1734 cm^-1^ for C=O stretch, 1627 cm^-1^ for N-H bend, 1377 cm^-1^ for sp^3^ C-H bend, 1247 cm^-1^ and 1161 cm^-1^ for acyl C-O, 1027 cm^-1^ for alkoxy C-O and 873 cm^-1^ for alkene sp^2^ C-H bend. The comparison of neem leaves peaks at 3297 cm^-1^ for N-H_2_, 2927 cm^-1^, and 2857 cm-1 for C-H, 1733 cm^-1^ for C=O stretch, 1607 cm^-1^ for N-H bend, 1165 cm^-1^ for acyl C-O, 1107 cm^-1^ and 1037 cm^-1^ for alkoxy C-O with nQDs shows the shift of peaks in nQDs at 3367 cm^-1^, 1734 cm^-1^, 1627 cm^-1^, 1247 cm^-1^, 1161 cm^-1^ and 1027 cm^-1^, along with the formation of new bonds in nQDs at 1377 cm^-1^ and 873 cm^-1^.

The optical characteristic of nQDs is UV Fluorescence Spectroscopy. The fluorescence spectra of nQDs were recorded by using excitation wavelengths from 300nm to 600nm in water and ethanol as solvents. In ethanol, the fluorescence intensity at 672nm increased upon excitation wavelength from 300 to 400nm with a maximum at 400nm wavelength. The fluorescence intensity at 672nm decreased from 450nm to 600nm excitation wavelength, as shown in **Figure 3(a)**. The nQDs showed red fluorescence under UV light and green under visible light in ethanol, as shown in **Figure 3(c)**. In water, the emission peak at 678nm does not shift when the excitation wavelength changes. The emission peak was at 678nm, and the fluorescence intensity increased upon excitation from 350nm to 400nm, with maximum emission at 400nm, and then decreased upon excitation from 450nm to 600nm wavelength. The emission peak at 436nm upon excitation wavelength of 300nm, represents the maximum emission peak of nQDs. The nQDs showed other emission peaks for example at 453nm upon excitation with 350nm, 461nm upon excitation with 400nm, and 530nm upon excitation with 450nm, as shown in **Figure 3(b)**. The fluorescence intensity was 623.989 a.u. at 436nm, which decreased to 184.129 a.u. at 678nm. Therefore, the 678nm peak was overshadowed by the 436nm peak, causing the nQDs to appear green instead of red under UV light. The nQDs were green under UV light and transparent under visible light in water, as shown in **Figure 3(d)**.

**Figure 3.**
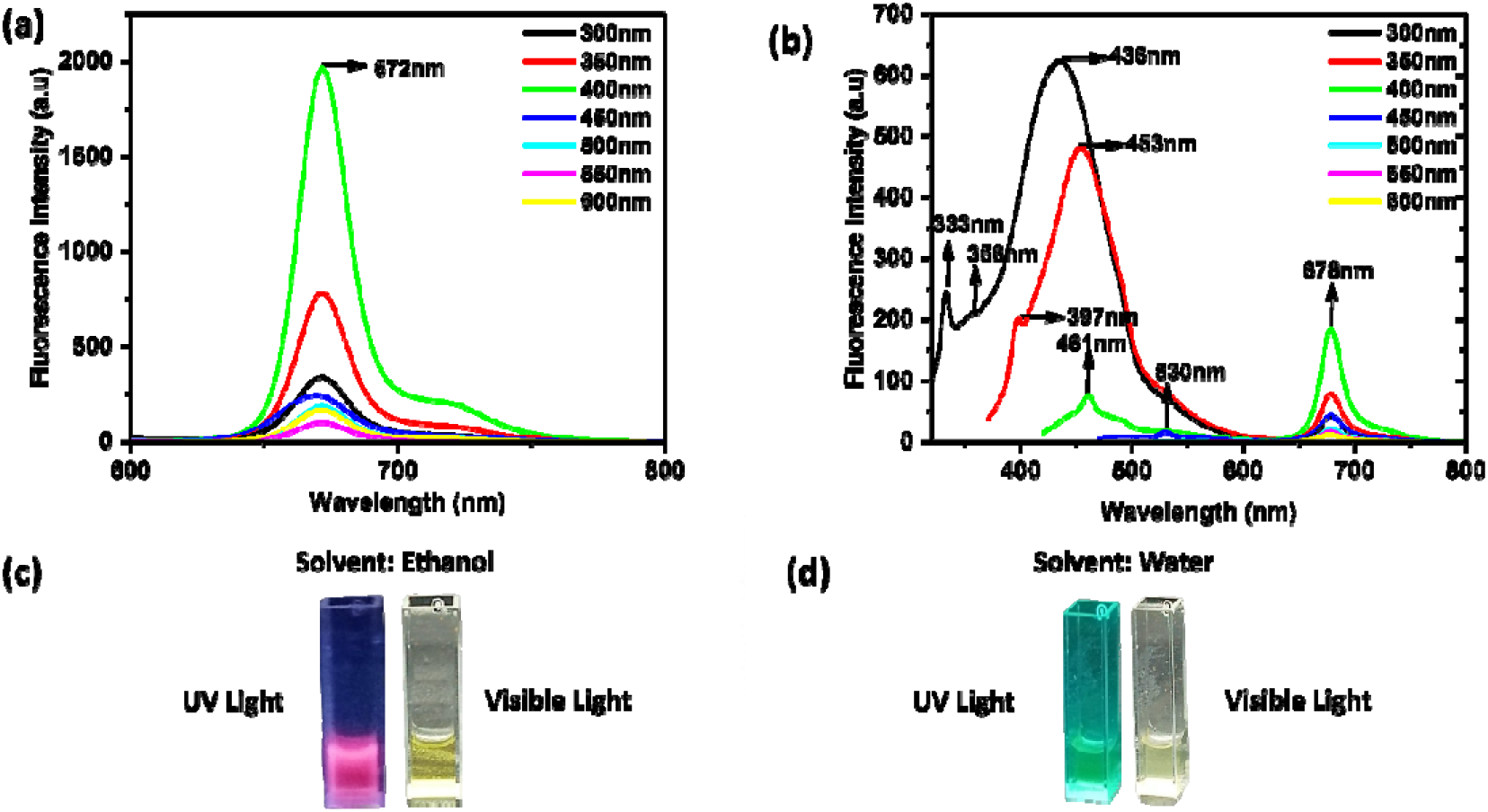
**(a)** At the 400nm excitation wavelength, the nQDs excitation and emission spectra reach their maximum at 672nm in ethanol. **(b)** Water was used as a solvent to dissolve nQDs. At the 300nm excitation wavelength, the nQDs excitation and emission spectra showed max at 436nm. Also, at the 400nm excitation wavelength, the nQDs’ excitation and emission spectra were demonstrated at 678nm. **(c)** The nQDs were red in UV light and green in visible light. (solvent-ethanol) **(d)** The nQDs were green in UV light and transparent in visible light. (solvent-water)

Atomic Force Microscopy (AFM) was used to study the quasi-spherical topography and morphology. With a scaled image, the particle size was measured using ImageJ software, and the Origin program was used to analyze the data. The nQDs were dissolved in ethanol and water, and then the size and height of the nQDs were measured. Firstly, water was used as the solvent, and nQDs showed their size and height at around 47nm and 1.16nm, respectively. **[Figure 4(a) & (b)]** Secondly, in ethanol as the solvent, nQDs size and height were measured at around 143nm and 24.8nm, respectively. **[Figure 4(c) & (d)]**

**Figure 4.**
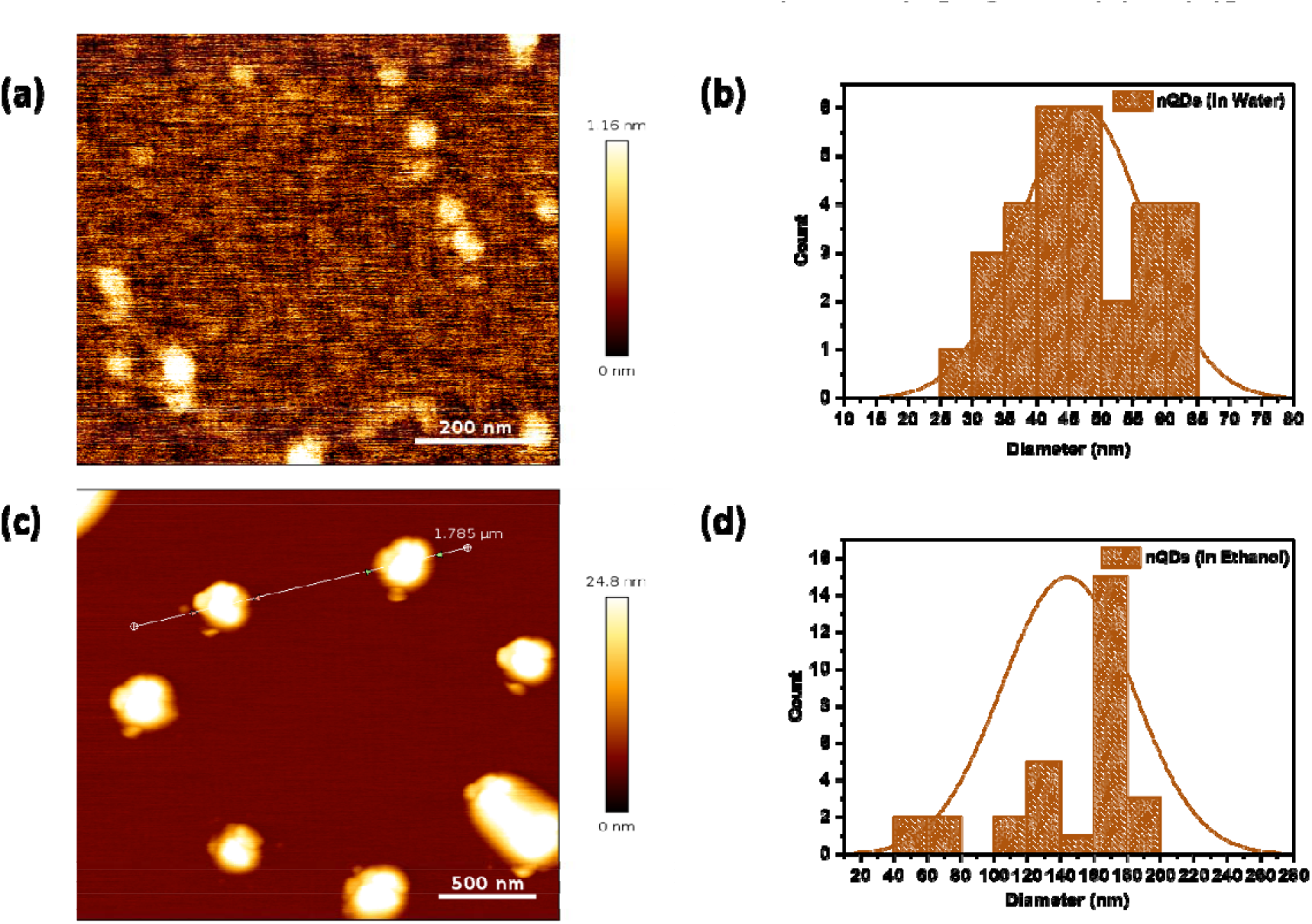
Atomic Force Microscopy (AFM): **(a) & (b)** AFM showed the nQDs particle size and height of approximately 47nm and 1.16 nm, respectively, in water. **(c) & (d)** AFM showed the nQDs particle size and height of approximately 143nm and 24.8nm, respectively, in ethanol.

### 3.2. Cellular uptake studies of nQDs

We used a 3-(4, 5-dimethyl thiazolyl-2)-2,5-diphenyltetrazolium bromide (MTT) to check the viability of the cells in the presence of nQDs. Two cell lines were selected: one is the retinal pigment epithelial (RPE1) cell line, and the other one is the breast cancer cells (SUM159A). A 96-well plate was seeded with approximately 1 10^5^ cells per well. We used a luminescence-based microplate reader (SpectraMax 190 Microplate Reader, Molecular Devices) to measure the results from both cell line tests. A range of increasing concentrations of nQDs (50, 100, 200, 300, 400, and 500 μg/mL) was added into RPE1 and SUM159A cells respectively. The percent viability was calculated following a 24-hour incubation period. In the SUM159A, the viability was 92%, 117%, 63%, 54%, 25%, and 9% at 50 100, 200, 300, 400, and 500 μg/mL concentrations respectively as shown in **Figure 5**. As the concentration of nQDs increases, the viability of the cells begins to decline. The observation of cancerous cells (SUM159A) demonstrates the anti-cancer characteristics of nQDs.

**Figure 5.**
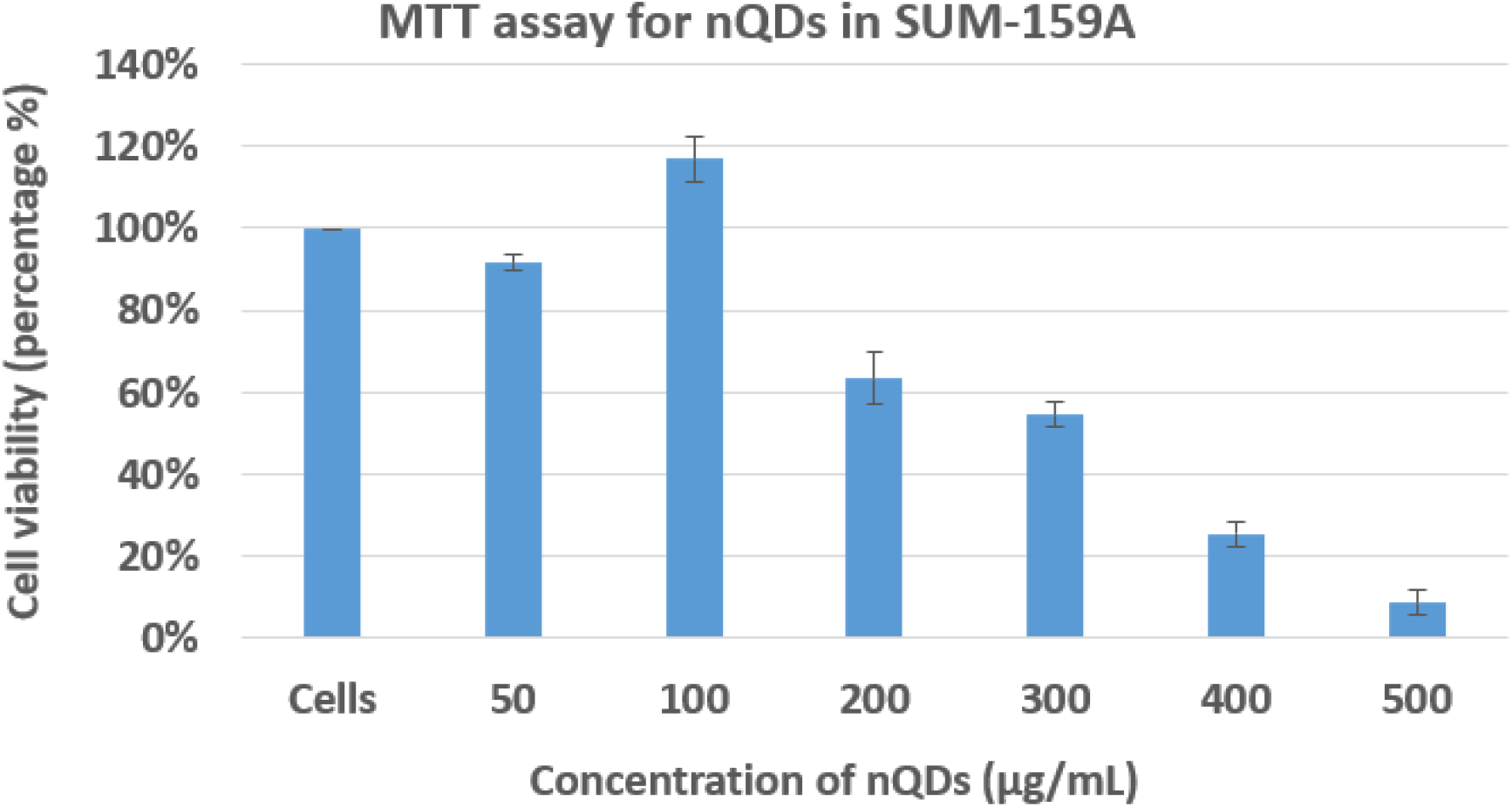
SUM159A cell viability at various concentrations of nQDs ranging from 0-500 μg/mL

To study potential use in bioimaging, the retinal epithelial pigment (RPE1) and SUM159A cells were cultured with nQDs at increasing concentrations and were analyzed using confocal microscopy analysis. The cells were treated with increasing concentrations of nQDs (50, 100, and 200 μg/mL) at 37 for 15 minutes demonstrating effective cell uptake of the nQDs by measuring their intracellular fluorescence.

In RPE Cells the fluorescence intensity increased with increasing concentration of nQDs demonstrating a concentration-dependent absorption and fluorescence from 50 to 200 μg/mL, as shown in **Figure 6**. However, in the case of SUM159A cells, the fluorescence intensity at 50 μg/mL showed maximum fluorescence, and at 100 μg/mL showed a decline in the fluorescence intensity again fluorescence intensity at 200 μg/mL increased with an increase in nQDs concentration. **Figure 7** demonstrates that the fluorescence intensity fluctuated with increasing nQDs concentration in SUM159A cells.

**Figure 6.**
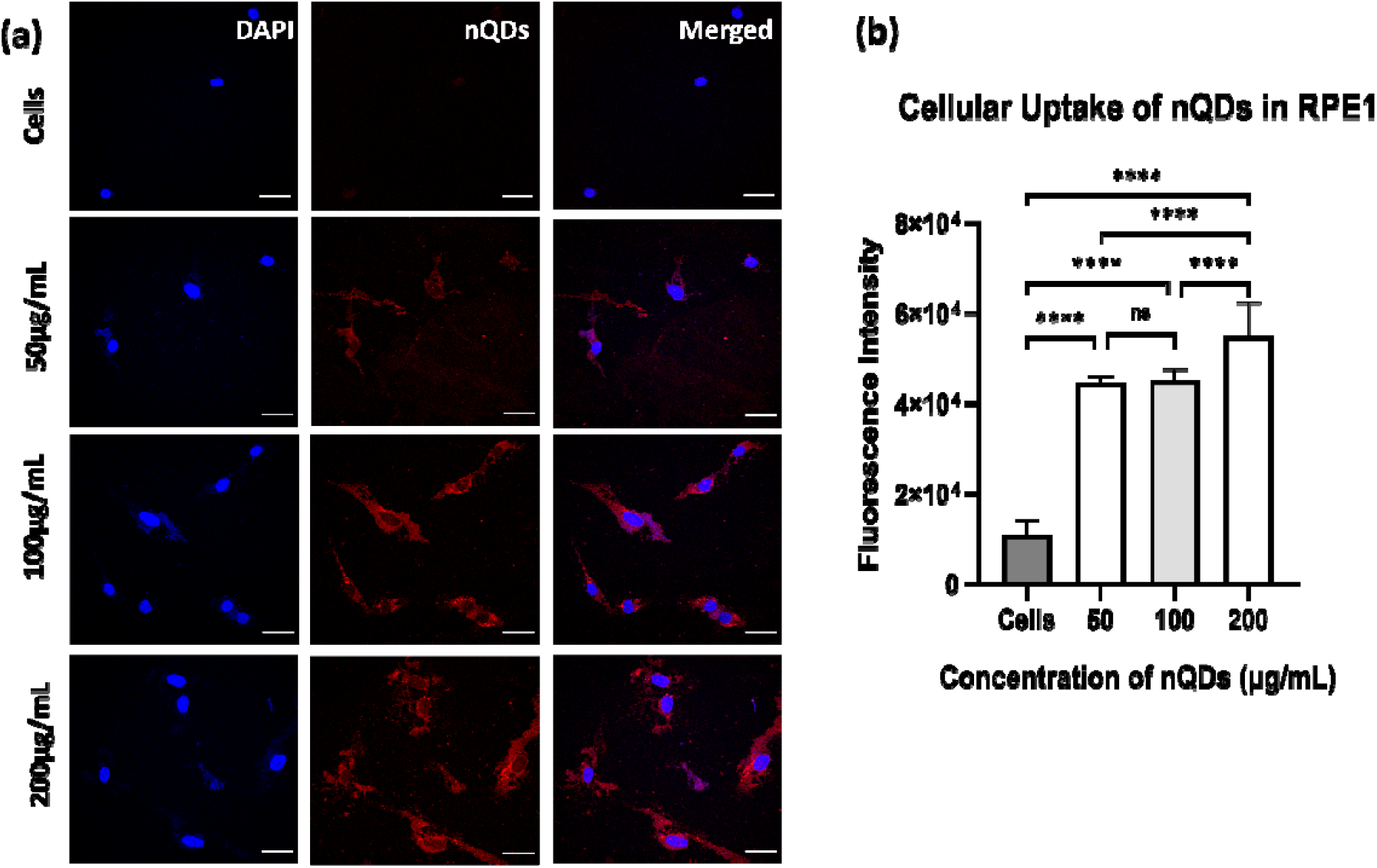
**(a) & (b)** In confocal microscopy images, Retinal pigment epithelial (RPE1) cells were treated with nQDs at 50, 100, and 200 μg/mL concentrations. A control without nQDs shows no fluorescence when excited at 633nm. Studies on the concentration-dependent characteristics of nQDs revealed that an increase in nQD concentration causes an increase in fluorescence intensity. The scale bar for confocal images is maintained at 5μm. The nQDs cellular uptake was also quantified at 50, 100, and 200μg/mL using Fiji ImajeJ Software. The statistical significance was tested by one-way ANOVA in the Prism Software and is represented as **** when p < 0.0001 and ns when there is no significant difference.(n=30)

**Figure 7.**
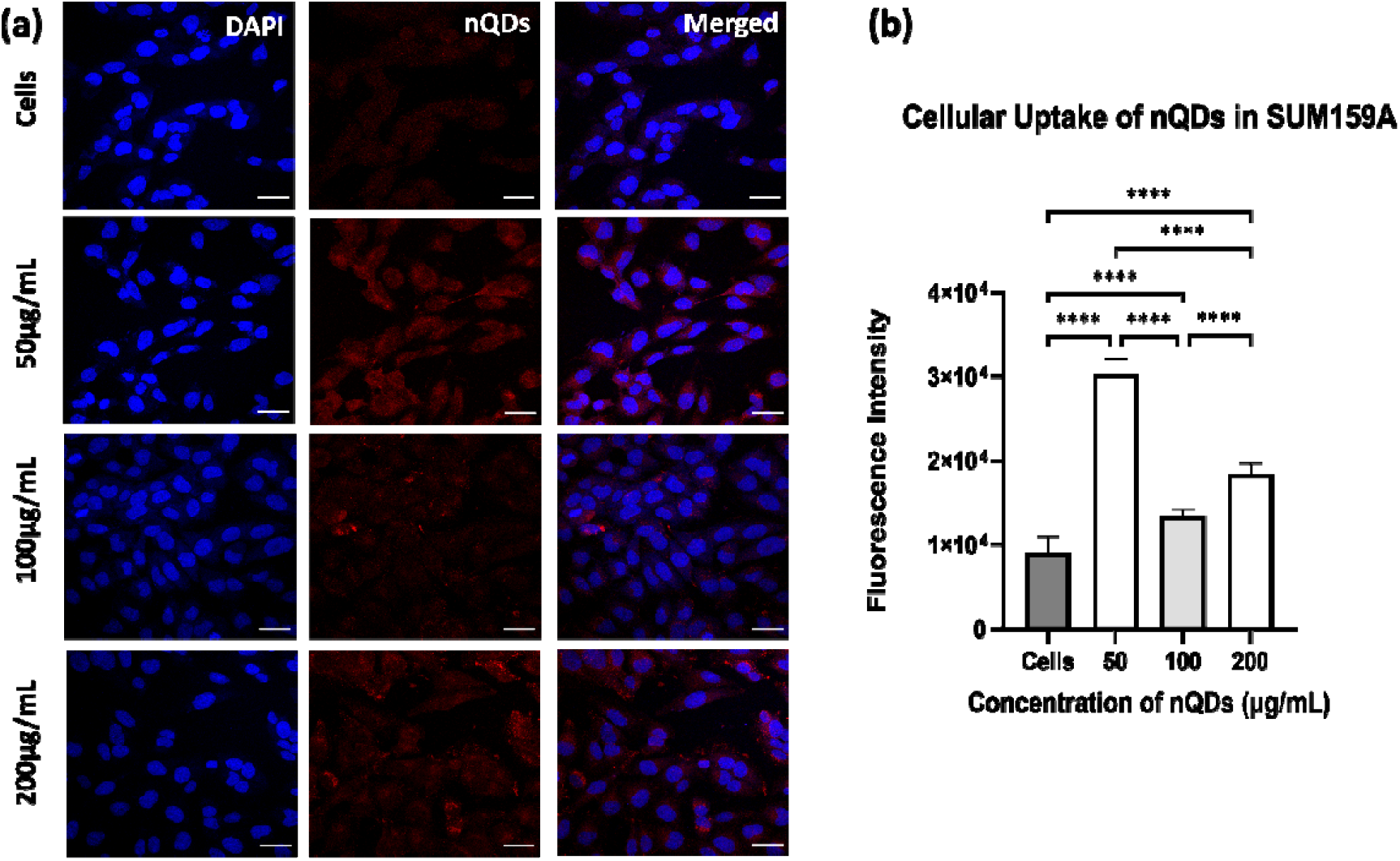
**(a) & (b)** SUM159A cells treated with nQDs at different concentrations of 50, 100, and 200 μg/mL are shown in confocal images. A control without nQDs shows no fluorescence when excited at 366nm. Studies on the concentration-dependent properties of nQDs revealed that an increase in nQDs concentration causes a fluctuation in fluorescence intensity. Fluorescence intensity at 50 μg/mL increases, 100 μg/mL reduces, and 200 μg/mL again rises. 5μm is the maintained scale bar at confocal images. The nQDs cellular uptake was also quantified at 50, 100, and 200 μg/mL using Fiji ImageJ Software. The statistical significance was tested by One-Way ANOVA in the Prism Software and is represented as **** when p < 0.0001. (n=30)

## 4. Discussion and Conclusions

*Azadirachta indica* (neem leaves) was used as precursor material and ethanol was used as a solvent. We synthesized red fluorescent carbon nanoparticles using a reflux method. The nQDs showed emission in red wavelength and had a lot of potential in biomedical applications in bioimaging. MTT assay showed nQDs biocompatibility in the case of non-cancerous cells and anti-cancer in the case of cancerous cells. Characterization techniques UV, fluorescence, FTIR, AFM, and DLS were used to analyze the chemical, physical, and optical properties of nQDs. The effect of solvents on fluorescence was studied in water and ethanol. A red color under UV light was shown in ethanol solvent. In ethanol, the fluorescence intensity was observed at 672nm at an excitation wavelength of 400nm. This intensity increased from 300 to 400nm upon and started to decrease from 450nm to 600nm upon excitation with different wavelengths, as shown in **Figure 3**. In water, the fluorescence intensity at 678nm was similar to that in ethanol. The nQDs were 43nm in size in water and in ethanol the nQDs aggregated to form 143nm size particles **Figure 4**. Furthermore, the nQDs cellular uptake study and biocompatibility assay showed anti-cancer properties in SUM159A cells. In addition, the cell viability also decreased with increasing the concentration of nQDs in SUM159A **Figure 5**. In the RPE1 cells, cellular uptake and fluorescence intensity increase with the increasing concentration of the nQDs. In RPE1 cells, the cellular uptake and fluorescence intensity increase with increasing nQD concentrations **Figure 6**. In the SUM159A cells, the cellular uptake and fluorescence intensity decreased with the increasing concentration of the nQDs. The fluorescence intensity was maximum at 50μg/mL concentration of nQDs and decreased at 100μg/mL and 200μg/mL **Figure 7**. Therefore, nQDs have shown excellent anti-cancer properties. Although the current work is still in its early stages, it gives us an essential fluorescent material that can be used and modified to enhance the optical properties and make it more functionalized and target-oriented for future biological applications such as bioimaging and anti-cancer applications.

## Acknowledgment

We sincerely thank all the members of the DB group for critically reading the manuscript and for their valuable feedback. CRTDH lab for fluorescence and UV spectra. CIF IITGN for confocal, AFM, DLS, and FTIR. SG is thankful to Sardar Patel University for doing research at IITGN. DB thanks SERB and Gol for the Core research grant, IITGN for the startup grant, and DBT-EMR, Gujcost-DST, and GSBTM for research grants. Imaging facilities of CIF at IIT Gandhinagar are acknowledged.

## References

1. Abid, N. et al. Synthesis of nanomaterials using various top-down and bottom-up approaches, influencing factors, advantages, and disadvantages: A review. Advances in Colloid and Interface Science vol. 300 Preprint at 10.1016/j.cis.2021.102597 (2022).

2. Thanh, N. T. K., Maclean, N. & Mahiddine, S. Mechanisms of nucleation and growth of nanoparticles in solution. Chemical Reviews vol. 114 7610–7630 Preprint at 10.1021/cr400544s (2014).

3. Pankhurst, Q. A., Thanh, N. T. K., Jones, S. K. & Dobson, J. Progress in applications of magnetic nanoparticles in biomedicine. J Phys D Appl Phys 42, 224001 (2009).

4. Titus, D., James Jebaseelan Samuel, E. & Roopan, S. M. Nanoparticle characterization techniques. in Green Synthesis, Characterization and Applications of Nanoparticles 303–319 (Elsevier, 2018). doi:10.1016/B978-0-08-102579-6.00012-5.

5. Quantum Dots. https://commons.wikimedia.org/wiki/File:Exicton_energy_levels.jpg#/media/File:Exicton_energy_levels.jpg.

6. Mohamed, W. A. A. et al. Quantum dots synthetization and future prospect applications. Nanotechnology Reviews vol. 10 1926–1940 Preprint at 10.1515/ntrev-2021-0118 (2021).

7. Barve, K., Singh, U., Yadav, P. & Bhatia, D. Carbon-based designer and programmable fluorescent quantum dots for targeted biological and biomedical applications. Materials Chemistry Frontiers Preprint at 10.1039/d2qm01287a (2023).

8. Brus, L. E. Electron-electron and electron-hole interactions in small semiconductor crystallites: The size dependence of the lowest excited electronic state. J Chem Phys 80, 4403–4409 (1984).

9. Ekimov, A. I., Efros, A. L. & Onushchenko, A. A. Quantum size effect in semiconductor microcrystals. Solid State Commun 56, 921–924 (1985).

10. Xu, X. et al. Electrophoretic analysis and purification of fluorescent single-walled carbon nanotube fragments. J Am Chem Soc 126, 12736–12737 (2004).

11. Lim, S. Y., Shen, W. & Gao, Z. Carbon quantum dots and their applications. Chemical Society Reviews vol. 44 362–381 Preprint at 10.1039/c4cs00269e (2015).

12. Pan, M. et al. Fluorescent carbon quantum dots-synthesis, functionalization and sensing application in food analysis. Nanomaterials 10, (2020).

13. Zhou, Y. et al. Multicolor carbon nanodots from food waste and their heavy metal ion detection application. RSC Adv 8, 23657–23662 (2018).

14. Xu, S. et al. Microwave-assisted synthesis of n, s-co-carbon dots as switch-on fluorescent sensor for rapid and sensitive detection of ascorbic acid in processed fruit juice. Analytical Sciences 36, 353–360 (2020).

15. Ding, H., Yu, S. B., Wei, J. S. & Xiong, H. M. Full-color light-emitting carbon dots with a surface-state-controlled luminescence mechanism. ACS Nano 10, 484–491 (2016).

16. Park, S. Y. et al. Photoluminescent green carbon nanodots from food-waste-derived sources: Large-scale synthesis, properties, and biomedical applications. ACS Appl Mater Interfaces 6, 3365–3370 (2014).

17. Mehta, S., Singh, U., Barve, K. & Bhatia, D. Psidium guajava derived carbon nanoparticles: A promising red emissive cellular bioimaging agent. doi:10.1101/2023.03.20.533411.

18. Wang, Y., Zhu, Y., Yu, S. & Jiang, C. Fluorescent carbon dots: Rational synthesis, tunable optical properties and analytical applications. RSC Advances vol. 7 40973–40989 Preprint at 10.1039/c7ra07573a (2017).

19. Cui, L., Ren, X., Sun, M., Liu, H. & Xia, L. Carbon dots: Synthesis, properties and applications. Nanomaterials vol. 11 Preprint at 10.3390/nano11123419 (2021).

20. Ge, G. et al. Carbon dots: Synthesis, properties and biomedical applications. Journal of Materials Chemistry B vol. 9 6553–6575 Preprint at 10.1039/d1tb01077h (2021).

21. Singh, P. et al. NON-TOXIC FABRICATION OF FLUORESCENT CARBON NANOPARTICLES FROM MEDICINAL PLANTS/SOURCES WITH THEIR ANTIOXIDANT ASSAY.

22. Sahu, S., Behera, B., Maiti, T. K. & Mohapatra, S. Simple one-step synthesis of highly luminescent carbon dots from orange juice: Application as excellent bio-imaging agents. Chemical Communications 48, 8835–8837 (2012).

23. Huang, H. et al. One-pot green synthesis of nitrogen-doped carbon nanoparticles as fluorescent probes for mercury ions. RSC Adv 3, 21691–21696 (2013).

24. Xu, Y. et al. Green synthesis of fluorescent carbon quantum dots for detection of Hg2+. Chinese Journal of Analytical Chemistry 42, 1252–1258 (2014).

25. Liu, W. et al. Green synthesis of carbon dots from rose-heart radish and application for Fe3+ detection and cell imaging. Sens Actuators B Chem 241, 190–198 (2017).

26. Jeong, C. J. et al. Fluorescent carbon nanoparticles derived from natural materials of mango fruit for bio-imaging probes. Nanoscale 6, 15196–15202 (2014).

27. Shehzadi, N., Sana, K., Masood, N., Afzal, U. & Ahmed Khan, M. A COMPREHENSIVE REVIEW: GREEN SYNTHESIS OF NANOPARTICLES FROM AZADIRACHTA INDICAPLANT. http://xisdxjxsu.asia.

28. Roy, P. et al. Plant leaf-derived graphene quantum dots and applications for white LEDs. New Journal of Chemistry 38, 4946–4951 (2014).

29. Meena, R. et al. Fluorescent carbon dots driven from ayurvedic medicinal plants for cancer cell imaging and phototherapy. Heliyon 5, (2019).

30. Yadav, P. K. et al. Green Synthesis of Fluorescent Carbon Quantum Dots from Azadirachta indicaLeaves and Their Peroxidase-Mimetic Activity for the Detection of H 2 O 2 and Ascorbic Acid in Common Fresh Fruits. ACS Biomater Sci Eng 5, 623–632 (2019).

31. Yadav, P. et al. Dopamine functionalized, red carbon quantum dots for in vivo bioimaging, cancer therapeutics, and neuronal differentiation. doi:10.1101/2023.06.16.545347.

